# Heterogeneity in polygenic scores for common human traits

**DOI:** 10.1101/106062

**Authors:** Erin B Ware, Lauren L Schmitz, Jessica Faul, Arianna Gard, Colter Mitchell, Jennifer A Smith, Wei Zhao, David Weir, Sharon LR Kardia

## Introduction

Genome-wide association studies (GWAS) on a wide range of important human traits have identified hundreds of variants in multi-cohort meta-analyses that have highly significant associations (p<5 x10^-8^) that replicate across studies. However, the identified genetic variants, or single nucleotide polymorphisms (SNPs), generally explain a very small fraction of variability in the trait of interest. Moreover, for most behavioral and health scientists, the use of such massive and complex genetic data is unwieldy. In response to the desire to harness genome-wide information, use of polygenic scores (PGSs), also known as “genetic risk scores”, has grown rapidly in the social, behavioral, and health sciences recently (see (1)).

PGSs use the results of GWAS—typically in the form of effect estimates or p-values for each nucleotide locus—to summarize an individual’s genetic association with a given trait. They can provide a single summary of measured genetic contribution of a trait that is easily integrated into more mainstream analyses of health and behavior. PGSs have multiple uses including improving risk prediction modeling controlling for some portion of variation due to genetics, investigating the common genetic basis among diseases, and estimating the genetic susceptibility of traits not measured but of high interest in a cohort (e.g. a PGS for post-traumatic stress disorder in a study of veterans or a cardiovascular PGS in a cognition study). PGSs are approximately normally distributed and relatively easy to construct. In short they appear to provide a relatively straightforward mechanism to integrate large amounts of genetic information into studies that are too small for genetic discovery, did not collect the health outcomes of interest, or do not have experience using large-scale genetic data.

The relative ease of creation and use has led to a wide range of practices in creating PGSs (2–4). These practices vary based on researchers’ decisions, including: 1) using genotyped SNPs versus imputed SNPs, 2) selecting SNPs for inclusion in the PGS from a meta-analysis of GWAS studies based on a particular p-value threshold, 3) whether to account for linkage disequilibrium (LD) across the genome, and 4) the effect of different options for accounting for linkage disequilibrium. And yet, despite their rapid adoption in the social sciences, best practices for PGS construction and evaluation remains relatively unexplored. Many researchers use programs such as PLINK (5), PRSice (6), and LDPred (7), designed to make PGS creation relatively simple. For novice users, these programs have produced a ‘black box’ creation platform that does not explore and display the range of PGSs that could be estimated. Moreover, researchers rarely report in sufficient detail the decisions, thresholds, and options used in the construction of their scores, which is likely to affect the replicability and reliability of PGSs. The degree to which all of these decisions and the combination of these decisions influence the performance of the PGSs is generally unknown, which may help explain contradicting results of the effectiveness of PGS in health and behavioral research.

Using the Health and Retirement Study (8) (HRS) we examine PGSs and their bootstrapped distributions for four traits height (stdHeight), body mass index (BMI), educational attainment (with SNP weights from two separate GWAS performed in 2013 (EA 2013) and 2016 (EA 2016)), and depression (MDD)) estimated using a) seven different p-value thresholds applied to the GWAS meta-analysis for SNP inclusion in the PGS (pT), b) genotyped versus imputed SNPs, and c) not accounting for linkage disequilibrium versus accounting for linkage disequilibrium using clumping or pruning for d) two different ancestry groups (non-Hispanic Black and non-Hispanic White). **Table 1** details the 434 distinct PGS estimation approaches (24 with no LD trimming + 216 with LD clumping + 192 with LD pruning + 2 top SNPs) we investigated using the SNPs and effect estimates from large published GWAS meta-analysis results where HRS was not present or was removed (9–13). To better understand the implications of the different options that underlie PGS creation we estimated how much of the phenotypic variation each score explained using semi-partial-R^2^ (with 95% bootstrapped confidence intervals), and the correlations among HRS participants’ PGSs within a trait.

**Table 1.**
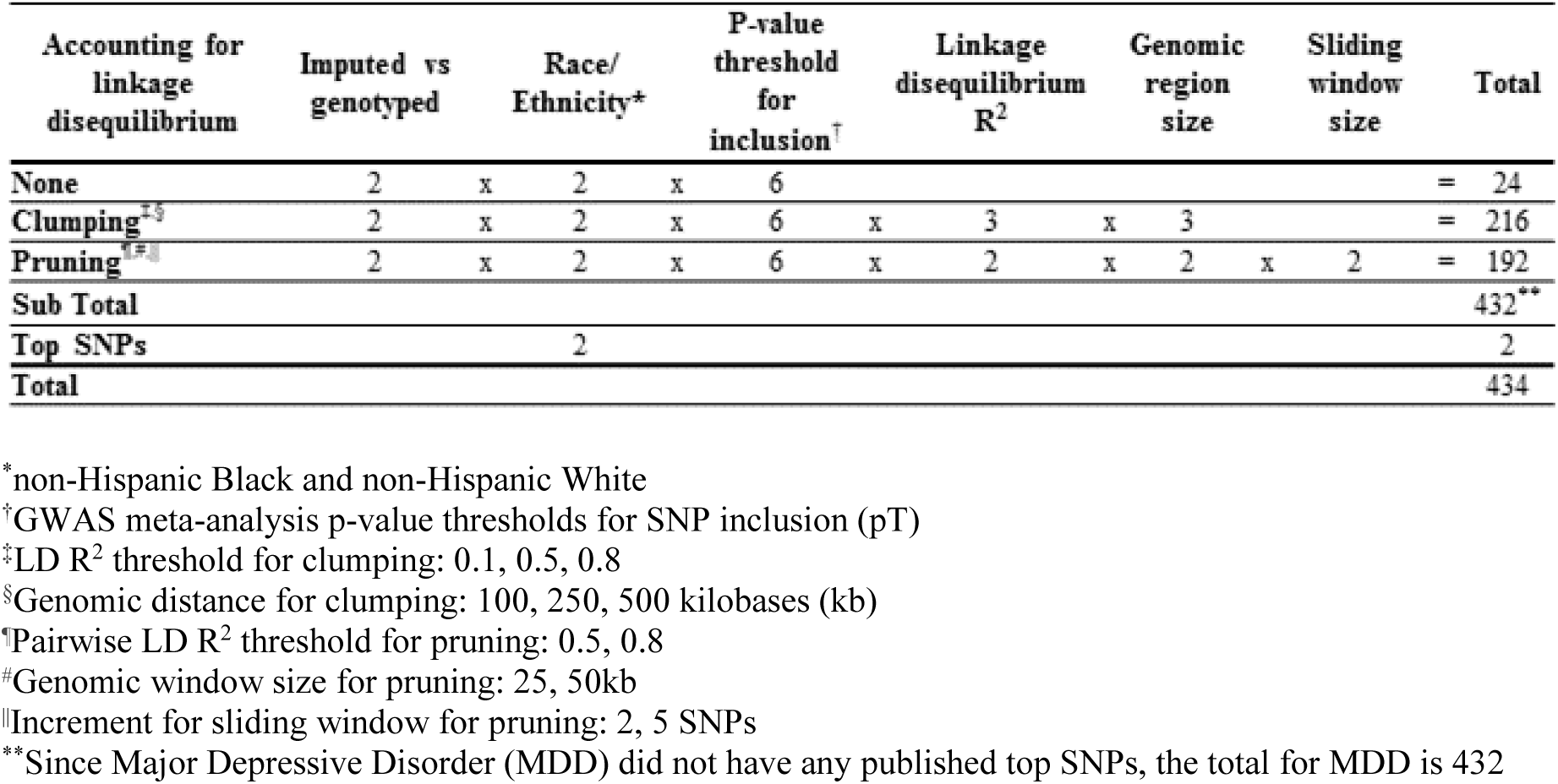
**Estimation approaches for polygenic score creation for each trait**

## Results

Given the complexity of PGS estimation procedures, we focused on examining the impact of using genotyped versus imputed SNPs, different p-value thresholds, and different LD trimming methods on the predictive capacity and correlation among scores within and among race/ethnicity.

### Directly genotyped SNPs versus imputed SNPs

In genomic studies, imputation allows for researchers to accurately estimate the genotypes of many unmeasured SNPs when a sample’s SNPs are paired with a well-established reference panel of more densely genotyped or sequenced individuals (e.g. HapMap, 1000 Genomes, Haplotype Reference Consortia (HRC)). Imputed SNPs are routinely used in GWAS to substantially increase the number of overlapping SNPs across studies and often genome-wide significant SNPs are found only in imputed data. Thus using imputed data may provide more overlapping SNPs between a study and the GWAS results, potentially improving the score. However, we found that there does not appear to be a significant difference between genotyped and imputed semi-partial R^2^, except in some isolated cases. There were a total of 108 paired comparisons per trait: 6 for no LD trimming (representing six different pTs), 54 for LD clumping (6 pTs x 3 LD R^2^ x 3 region sizes), and 48 for LD pruning (6 pTs x 3 LD R^2^ x 3 region sizes x 2 sliding window options). In **Table 2** we compare the percent variation explained by PGSs estimated using genotyped versus imputed data in ethnic groups separately. Examining results by LD trimming method reveals some interesting patterns. First, with no LD trimming, for the majority of the paired comparisons, observed R^2^_genotype_ is greater than the corresponding observed R^2^_imputed_. In fact, 95% confidence intervals around genotyped and imputed R^2^s did not overlap for five of these cases (see **Table S1**, **Fig. S1**), indicating statistically significant differences in the amount of trait-variation explained in this part of the PGS estimation space. All five of these PGS pairs were estimated in non-Hispanic White, for the EA 2016 trait. In contrast, when using the LD clumping method, genotyped SNPs outperformed imputed SNPs 58.7% (317/540) of the time and, when using LD pruning, the imputed SNPs appeared to be better than genotyped in 52.9% (226/480) of the paired comparison. Overall, by comparing 95% confidence intervals, we found no paired comparisons where the R^2^_imputed_ was significantly higher than the R^2^_genotype_ (**Table 2, Fig. S2/Fig. S3**).

**Table 2.** **Comparing polygenic scores from directly genotyped SNPs to imputed SNPs across estimation approaches**

| Race/ethnicity | Trait | Linkage disequilibrium (LD) trimming method |  |  |
| --- | --- | --- | --- | --- |
|  |  | None (m*=6) | Clumping (m*=54) | Pruning (m*=48) |
| | | $\uparrow R^2_{\text{genotype}} > R^2_{\text{imputed}}$<br>ed<br>n<sup>‡</sup> (%) | $\uparrow R^2_{\text{genotype}} > R^2_{\text{imputed}}$<br>ed<br>n<sup>‡</sup> (%) | $\uparrow R^2_{\text{genotype}} > R^2_{\text{imputed}}$<br>ed<br>n<sup>‡</sup> (%) |
| Non-Hispanic Black | Body mass index | 2 (33.3) | 15 (27.8) | 8 (16.7) |
|  | Educational attainment 2013<sup>†</sup> | 5 (83.3) | 18 (33.3) | 11 (22.9) |
|  | Educational attainment 2016<sup>§</sup> | 4 (66.7) | 47 (87) | 46 (95.8) |
|  | Sex standardized and adjusted height | 3 (50) | 28 (51.9) | 9 (18.8) |
|  | Major Depressive Disorder | 5 (83.3) | 45 (83.3) | 30 (62.5) |
| Non-Hispanic White | Body mass index | 6 (100) | 27 (50) | 24 (50) |
|  | Educational attainment 2013<sup>§</sup> | 5 (83.3) | 24 (44.4) | 11 (22.9) |
|  | Educational attainment 2016<sup>¶</sup> | 6 (100)<sup>#</sup> | 48 (88.9) | 48 (100) |
|  | Sex standardized and adjusted height | 6 (100) | 35 (64.8) | 23 (47.9) |
|  | Major Depressive Disorder | 4 (66.7) | 30 (55.6) | 16 (33.3) |

\*m is the number of paired comparisons between semi-partial $R^2$ for the polygenic score created from directly genotyped versus imputed, within race/ethnicity
<sup>†</sup> $R^2_{\text{genotype}}$ is the semi-partial $R^2$ for the regression between the polygenic score and trait, adjusting for covariates when directly genotyped SNPs were used, $R^2_{\text{imputed}}$ is the semi-partial $R^2$ for the regression between the polygenic score and trait, adjusting for covariates when imputed SNPs were used
<sup>‡</sup>n is the number of paired comparisons between semi-partial $R^2$ for the polygenic score created from directly genotyped versus imputed, where the mean $R^2_{\text{genotype}}$ is greater than the mean $R^2_{\text{imputed}}$
<sup>§</sup>2013 uses SNP weights from the Educational attainment GWAS published in 2013(11)
<sup>¶</sup>2016 uses SNP weights from the Educational attainment GWAS published in 2016(12)
<sup>#</sup> Five of the six paired comparison for Educational attainment using 2016 GWAS weights and no LD trimming had $R^2_{\text{genotype}}$ significantly greater than $R^2_{\text{imputed}}$

Given the stochastic nature of clumping and pruning, the HRS has decided to create PGSs using genotyped data and no LD trimming at a p-value threshold for SNP inclusion of 1 (pT=1) to enable replication by other researchers. In **Fig 1**, we describe both the observed semi-partial R^2^ and the 95% bootstrapped confidence intervals for each of the traits in both race/ethnicities for these specifications.

**Figure 1.**
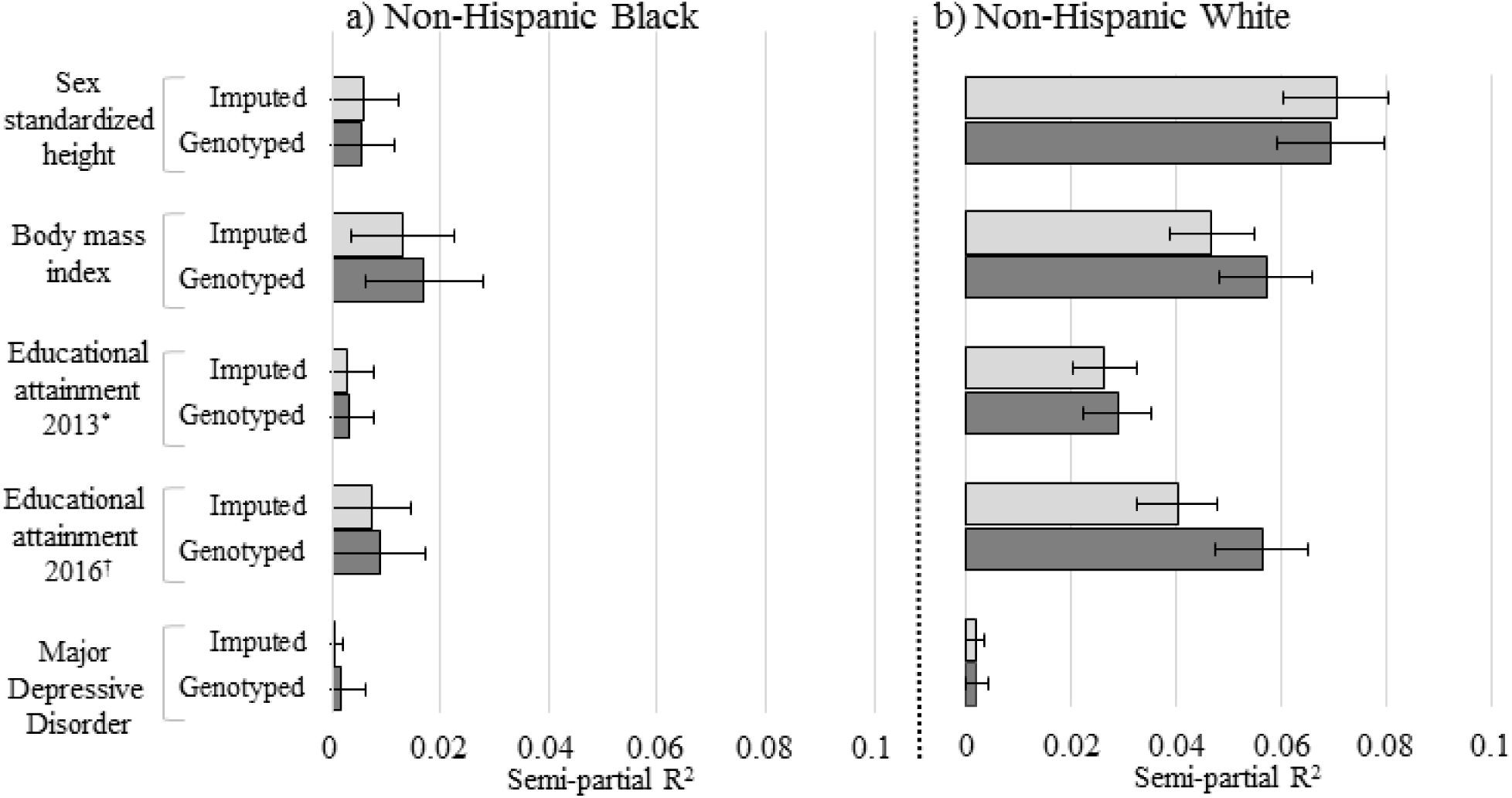
Comparison of semi-partial R^2^ for polygenic scores created from directly genotyped and imputed SNPs, by ethnicity and trait. No LD trimming, pT = 1.0. BMI: body mass index, EA: educational attainment, MDD: Major Depressive Disorder. Semi-partial R^2^ and 95% confidence intervals from bootstrapped models with 1,000 repetitions. Models include covariates reflective of the meta-analysis GWAS from which the SNP weights came.

To further examine the implications of using genotyped vs. imputed SNPs, we investigated the correlations between HRS participant’s PGSs that were estimated through different approaches. In **
Fig. 2**, each box plot represents the distribution of Pearson’s correlation coefficients between 18 different estimation methods (nine clumping, eight pruning, and one with no LD trimming) of PGSs within a pT set. The PGSs created using directly genotyped data (dark gray in **Fig. 2**) tended to have lower median correlations and wider distributions compared to imputed PGSs (light gray) within a trait for a given pT **(e.g.** Non-Hispanic Whites stdHeight at pT=0.001: median r_genotyped_ = 0.83, median r_imputed_ = 0.92, **Fig. 2)**. These patterns are consistent even where the base GWAS meta-analysis results have been heavily trimmed for LD (whether pruned or clumped), as in the case of the PGSs created using effect estimates from the Psychiatric Genomics Consortium MDD results (see **supplemental methods S3, Fig. S2.d, S2.d, Fig. S4 Panel 5, and Table S1.e**). Within the traits evaluated, these aforementioned patterns are consistent across race/ethnicity. For traits that have reported replicated, independent genome-wide significant (p-value < 5x10^-8^) SNPs (i.e. top SNPs), the box plot representing the correlation between that top SNPs score and all of the other estimation methods across all pTs appears first. Since there is only a single PGS that can be estimated from the top genome-wide significant SNPs reported in the meta-analysis, we estimated the HRS participants’ correlations with all other 108 PGSs estimated (6 no LD trimming + 54 LD clumping + 48 LD pruning). In all cases, the PGS from the top SNPs are poorly correlated with other PGSs.

**Figure 2.**
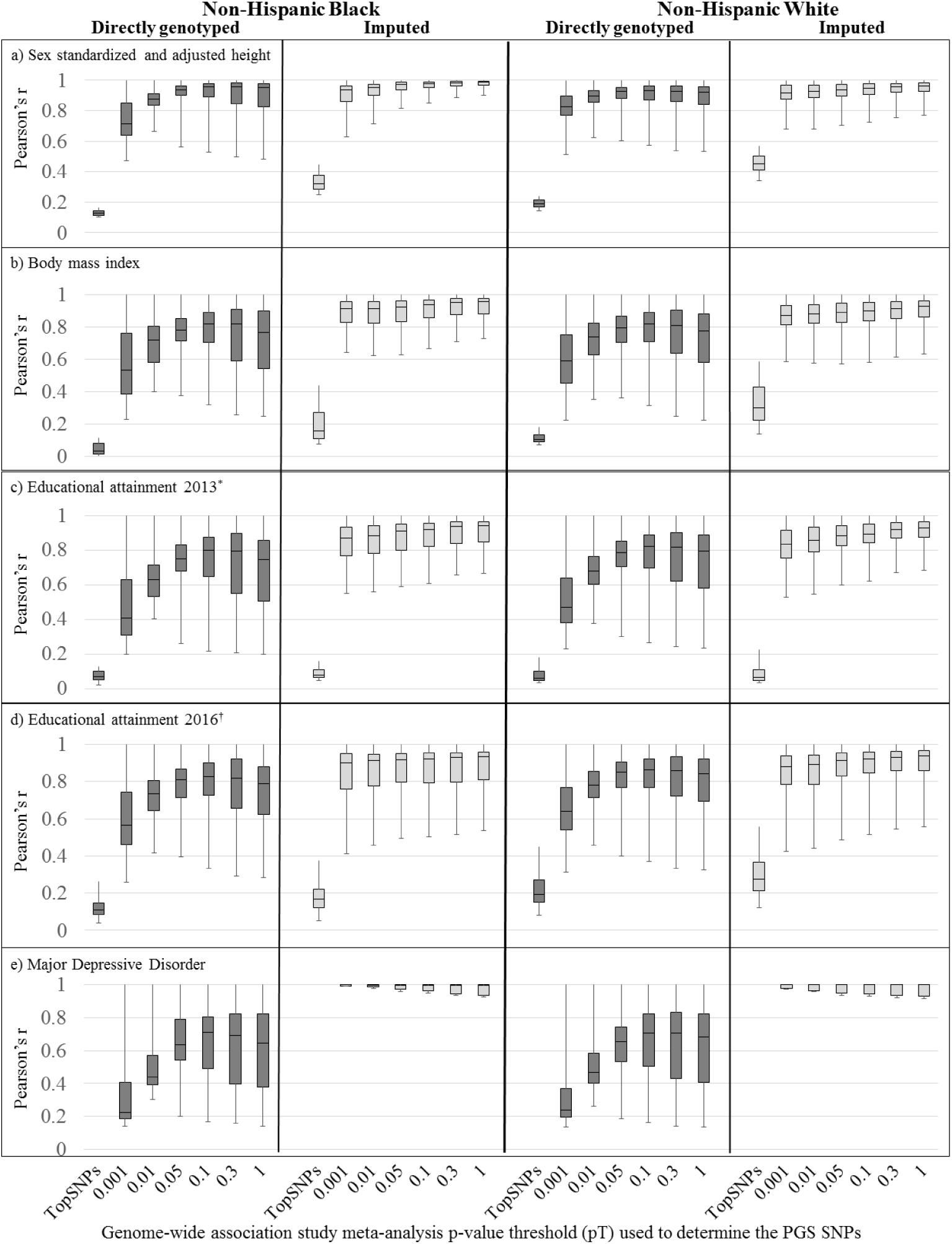
Distributions of Pearson’s correlation coefficient among HRS participant polygenic scores from different estimation approaches, by p-value inclusion threshold. Distributions of Pearson’s correlation coefficient among HRS participant polygenic scores from different estimation approaches, by p-value inclusion threshold. **Dark gray:** correlations between 18 different LD methods at six different p-value threshold cutoffs (plus a top SNP correlation set that includes all other 108 estimation approaches) for PGSs created with directly genotyped SNPs. **Light gray:** correlations between 18 different LD methods (1 no LD trimming + 8 LD clumping + 9 LD pruning) at six different p-value threshold cutoffs (plus a top SNP correlation set that includes all other 108 estimation approaches) for PGSs created with imputed SNPs. *Top SNP correlation set includes all other 108 estimation approaches. There were no genome-wide significant SNPs from the Major Depressive Disorder GWAS. ^†^2013 uses SNP weights from the Educational attainment GWAS published in 2013 (11) ^‡^2016 uses SNP weights from the Educational attainment GWAS published in 2016 (12)

### P-value threshold

When p-value thresholds of 1.0, 0.3, 0.1, 0.05, 0.01, 0.001, and 5×10^-8^ (top SNPs representing a genome-wide significant hits) are applied to the GWAS meta-analysis results, they coincide with progressively smaller numbers of SNPs passing into the set of SNPs available to enter the PGS calculation **(Table S1)**. Each p-value threshold represents a different balance in including the SNPs with the strongest signal vs larger coverage of the whole genome, which may also include more random noise. The subset of top SNPs (i.e. those that were genome-wide significant in the GWAS meta-analysis) for each trait predicted the lowest amount of trait-variation, or very close to it, across the estimation space (BMI: **Fig. 3,** all other traits: **Figs. S2, S3)**. While the estimation space is complex, we observed that PGSs estimated from directly genotyped SNPs at the lower GWAS meta-analysis p-value thresholds (pT=0.001 and 0.01) for BMI, EA 2013, EA 2016, and MDD often explained less of the percent trait-variation compared to higher pT thresholds (pT=0.05, 0.1, 0.3, 1.0). In non-Hispanic Blacks, the PGS from the higher pTs are the only non-zero PGSs identified – independent of whether or not directly genotyped or imputed SNPs were used. Overall, the effects of pT and LD trimming appear to be very consistent between genotyped and imputed PGSs.

**Figure 3.**
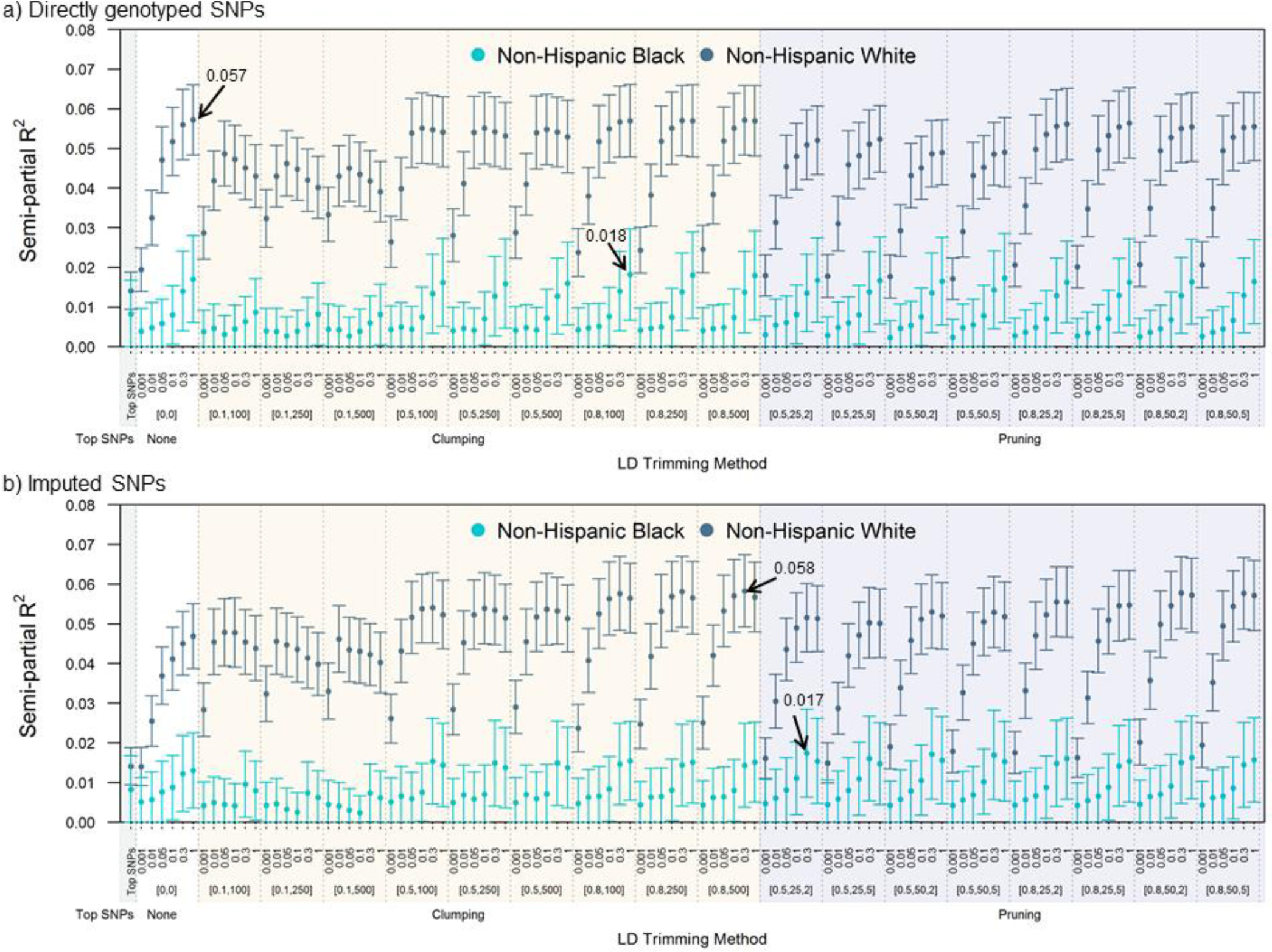
Semi-partial R^2^ with 95% standard error bars for regression of body mass index onto the polygenic score. **a)** directly genotyped and **b)** imputed SNPs. Arrows represent the highest amount of observed trait-variation explained for each race/ethnicity. Semi-partial R^2^ and 95% standard errors were created from 1,000 bootstrapped repetitions, adjusting for covariates that were used in the GWAS meta-analysis from which the SNP weights came. BMI: body mass index, LD: Linkage Disequilibrium. None: no LD trimming; LD Pruning [LD R^2^, window size (kb), sliding increment (number of SNPs)], LD Clumping [LD R^2^, region size (kb)]

A notable exception to these trends was the PGSs estimated for sex-standardized and adjusted height, where PGSs constructed with lower pT had higher percent trait-variation explained (**Figs. S2, S5**). For example, using directly genotyped SNPs, the PGS with the highest percent trait-variation explained in the non-Hispanic Whites was 10.4% (clumping LD R^2^ 0.1, region size 500 kb, pT=0.001, 95% bootstrapped CI: 9.2, 11.6). This estimation approach (clumping with an LD R^2^=0.1) was consistently in the set of lowest performers across the other traits. In contrast, the other estimation approaches using the pT=0.001 produced PGSs that often had significantly lower R^2^ than the top performers (non-Hispanic White: stdHeight, BMI, EA 2013, EA 2016; **Fig. S5**).

Examining the distribution of correlations among the HRS participant’s PGSs (**Fig. 2**) from the different estimation methods as a measure of the robustness across the estimation space, conditional upon the pT, we see wide variability in correlations across estimation approaches. The PGSs created using SNPs from the lower GWAS meta-analysis pTs (pT=0.001 and 0.01) had lower median Pearson’s correlation than those at higher pTs (pT=0.05, 0.1, 0.3, 1.0, **Fig. 2)**. Similar to what we observed in **Fig. S5** with the numerous scores that are not significantly different than the top performer (highest semi-partial R^2^), we note that the higher pTs gave rise to a higher level of consistency across all estimation methods for imputed PGSs as shown in the boxplots of the correlations in **Fig. 2** compared to the pTs at 0.001 and 0.01. This was not the case for genotyped scores. In particular, the top SNPs scores consistently had the lowest correlations with other methods across the estimation space. Looking across the entire estimation space in **Fig. 4**, a checkerboard pattern illustrates the lack of correlation between PGSs estimated with pT=0.001 relative to the other pTs (BMI: **Fig. 4**, all other traits **Fig. S4**).

**Figure 4.**
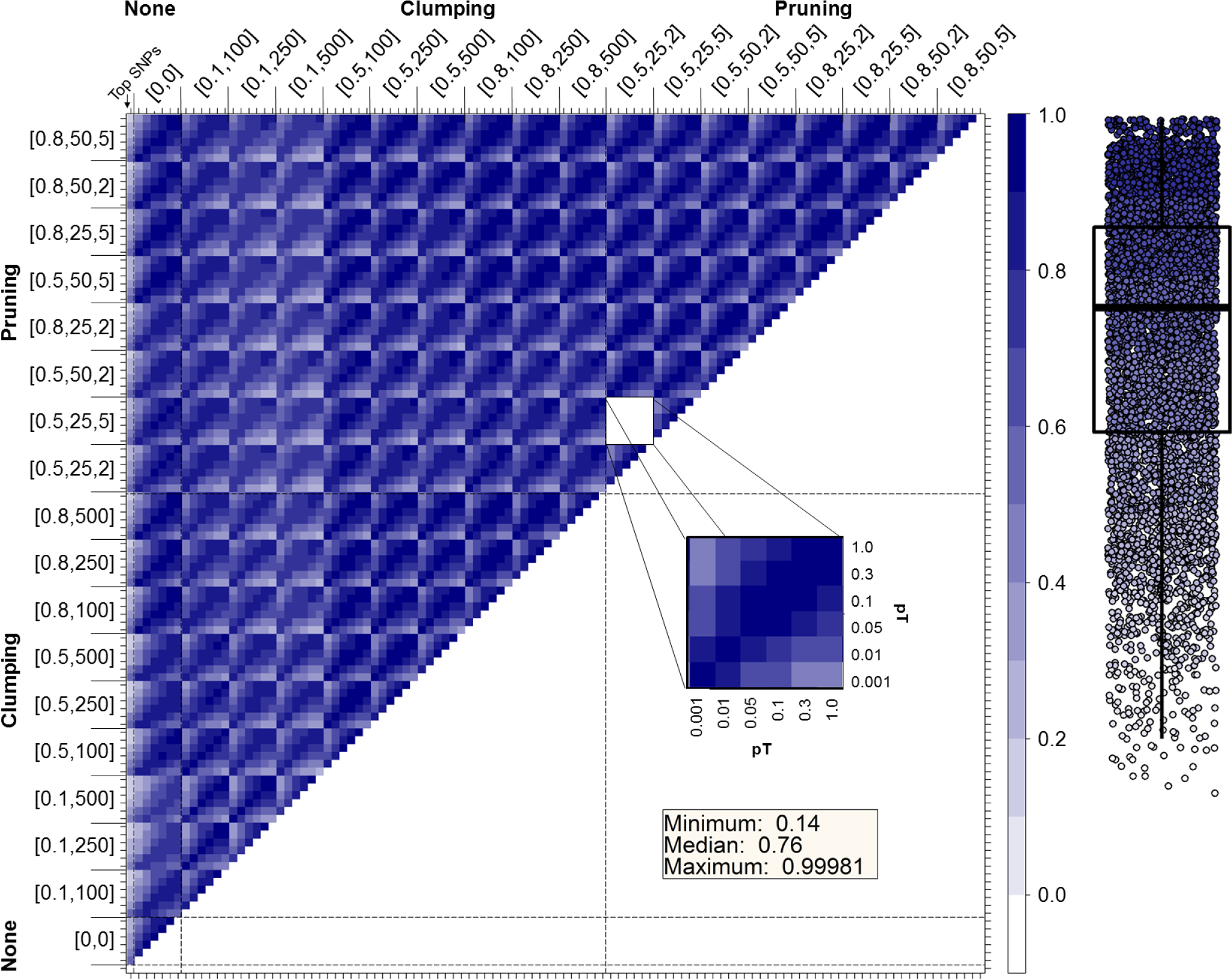
Pearson’s correlation between each of 109 polygenic scores for body mass index. Pearson’s correlation coefficients between each of 109 polygenic scores for body mass index, by ethnicity, shaded from −0.1 (light) to 1.0 (dark) using directly genotyped SNPs. BMI: body mass index, LD: Linkage Disequilibrium. None: no LD trimming; LD Pruning [LD R^2^, window size (kb), sliding increment (number of SNPs)], LD Clumping [LD R^2^, region size (kb)]. Inner boxes represent different GWAS meta-analysis p-value thresholds (pTs) (see inset)

### LD trimming and LD trimming options

In **Fig. 3** we display the range of semi-partial R^2^ values and corresponding bootstrapped 95% confidence intervals, for the regression of BMI on PGSs calculated using each estimation method at each pT, separating the results by LD trimming methods (top SNPs, none, clumping, pruning). The type of LD pruning appears to have much larger effects on the PGS estimation than the choice of genotyped versus imputed SNPs – that is, there are many equal results, even though they include different SNPs. Overall, there are no clear winners for which LD trimming method parameters produce the best PGS that explains largest trait-variation. It does appear that greater inclusion of SNPs results in higher R^2^s (**Fig. 3**). This finding is somewhat different for stdHeight (**Fig. S5**). In the non-Hispanic Whites, the clumping methods with LD R^2^=0.1 tended to have a dip in performance compared to the other clumping and pruning methods for BMI, EA 2013, and EA 2016, but not stdHeight (**Fig. 3, Figs. S1, S3**). The majority of the PGS estimates for MDD were non-significant predictors of the trait-variation in either non-Hispanic Whites or non-Hispanic Blacks and do not provide any insight into the LD trimming impact on PGS estimation.

### Cross-race/ethnicity comparison

For BMI, EA 2013, EA 2016, and stdHeight, a majority of the PGSs predicted significantly more trait variation in non-Hispanic Whites than non-Hispanic Blacks, for each estimation approach (**Fig 3, Fig. S3**). These differences are most likely due to factors outside of the PGS estimation, such as mutational spectra represented in the genotyping chips, differences in the race/ethnic origin of the cohorts included in the GWAS meta-analysis, and differences in the etiology of the phenotype between race/ethnicity (i.e. educational attainment).

## Discussion

This work demonstrates that estimation decisions for creating PGSs may have high heterogeneity, less than optimal predictive power, and can give rise to inaccurate inferences about the role of genetic variation in the trait variation across PGSs even within the same study.

Previous evaluation of PGS estimation methods have focused on predictive power, varying several of the decisions involved in the process. Several studies have investigated the effect of the distributional assumption imposed on the SNPs in the GWAS on the predictive power, concluding that the sample size for these GWAS needs to be upwards of 10,000 cases and controls for diseases controlled by 1000 loci with a mean relative risk of 1.04 (14, 15) and hundreds of thousands for SNP weights to be valid for risk prediction in more complex traits (16). Limiting the SNPs that enter the PGS through selection of only functional SNPs, adjusting the GWAS weights for “winners curse”, or inferring posterior mean effect sizes for each marker have shown limited success in boosting the predictive power of the PGS (7, 17).

Others have found, consistent with our work, that the pT has a substantial effect on the performance of PGSs. However, which pT is the optimal one varies by trait. Two studies found that prediction is optimized by selecting variants with pT as high as 0.001 (18, 19) – though these thresholds were optimal only for the trait or type of trait analyzed (binary trait with prevalence of 10% (18), coronary heart disease (19). We have found the landscape to be much more complex, and the optimal pT to be trait-dependent. Our work extends these discoveries by examining multiple attributes of the estimation process simultaneously to better elucidate the combined effects of different decisions in the estimation space on not only the predictive power of the PGS, but the distributions and relationships between PGS.

Overall, we have several key findings from this work. First, PGSs created using approaches decisions may explain significantly different amounts of variation in the phenotype of interest (see **Fig. S5**). Second, not only do PGSs estimated through different approaches explain different percentages of variability, but they also may represent potentially different genetic explanations, since the correlation between two scores was as low as r=0.02 (**Fig. S4, Panel 3**) for a given pT. Third, not all PGSs for a trait are necessarily significant predictors of their phenotype of interest. We found that 18.8% (205 out of 1088) of the PGSs in the non-Hispanic White sample were not significantly associated (α = 0.05) with the trait of interest. Fourth, PGSs created using weights derived from GWAS performed in individuals with European Ancestry generally do not perform as well in individuals with African Ancestry. In our study, the PGSs were significantly associated (p-value ≤ 0.05) with the traits in 81.2% (883 out of 1088) of the instances in non-Hispanic White but in only 29.4% of the non-Hispanic Black PGSs (320 out of 1088). At a more stringent significance criteria (α = 0.001), these percentages drop to 79.9% (869 out of 1088) and 5.2% (57 out of 1088), respectively. In general, the European Ancestry based weights produced PGSs in the HRS non-Hispanic Black sample that explained approximately half the amount of variability in the phenotype as the similarly weighted PGSs in the HRS non-Hispanic Whites. We did note, however, that the correlation structure between scores was similar in the non-Hispanic Black and White samples (**Fig. 2**, **Fig. S5**). Finally, though the “top SNPs” scores are the least computationally demanding, they consistently explained the least amount of trait-variation and also were not strongly correlated with PGSs created using other estimation methods (**Fig. 2**, **Figs. S2, S3, S5**).

While we attempted a thorough investigation of decisions that go into creating polygenic scores, there is no way to evaluate *all* decisions that influence the creation of a PGS. For instance, though we found that genotyped data work better for creating the PGSs, this result may be conditional on the genotyping chips that were used in the HRS (llumina HumanOmni2.5-4v1 and −8v1 BeadChips) and may not be true for studies using other genotyping chips. In addition, we were unable to fully evaluate all packages available for creating PGSs because of the lack of detailed GWAS meta-analysis results available from consortia that are required in these packages, particularly LDPred (7). Lastly, we were unable to evaluate the impact of using non-Hispanic Black GWAS findings on PGSs in HRS (both in how these weights would affect PGSs created for the non-Hispanic Black sample and how the cross-ethnic weighting would influence the scores created in the non-Hispanic White sample) because there are not yet large, replicated GWAS of all of these traits for non-Hispanic Blacks in the literature. While many trans-ethnic discovery findings suggest that biology does not greatly differ across ethnicities (20), the impact of using effect sizes of GWAS results from European Ancestry to build PGSs in other ethnicities has not been fully examined.

Based on our findings, and in light of the limitations of these analyses, we have highlighted several recommendations to researchers, as well as for consortia wishing to publish their meta-analysis results for public use (**Table 3**):

**Table 3.**
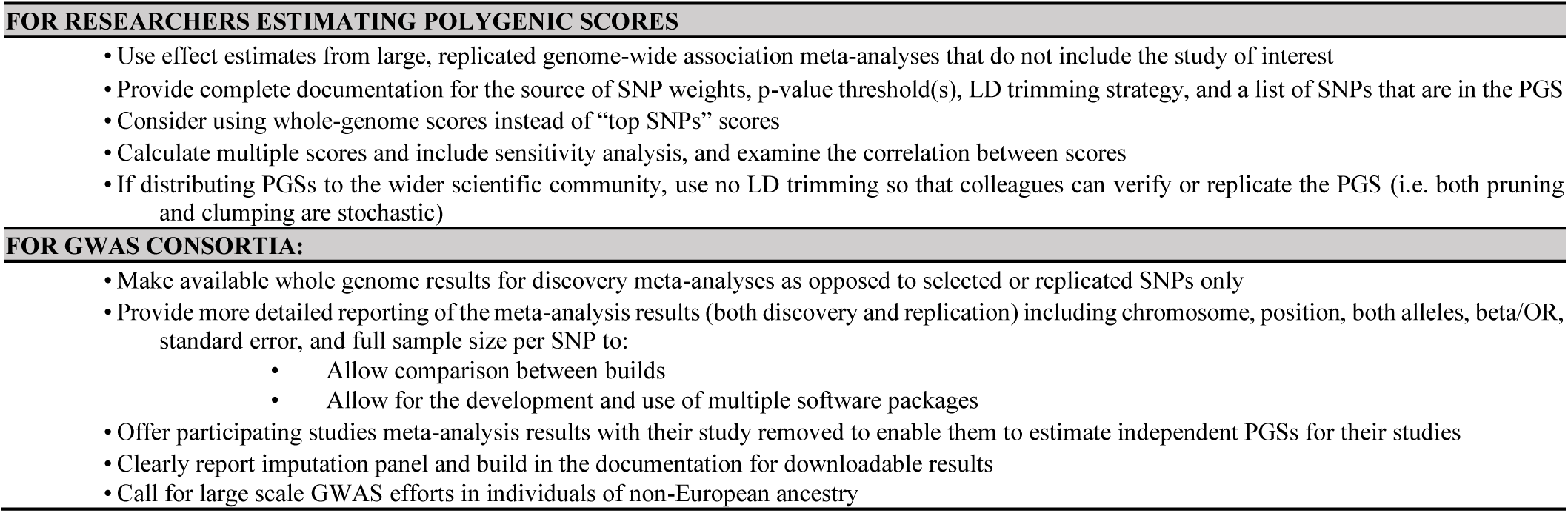
**Recommendations for researchers and GWAS consortia regarding polygenic score use**

Per these findings and recommendations, the HRS is creating PGSs for the wider scientific community using genotyped SNPs, no LD trimming, and a pT of 1 in order to enable replication by other scientists and to maximize the potential predictive capacity.

More work is needed to develop better and more consistent methods to estimate PGSs and incorporate them into a biosocial framework. PGSs expand our ability to integrate knowledge about an individual’s genetic susceptibility – a measure of their internal biological environment – into more complex and integrated biosocial modeling, even when the phenotype may not be measured explicitly. Never before have we had the ability to accurately represent individual-level genetic susceptibility for traits not measured explicitly (e.g. estimating Alzheimer’s genetic risk and incorporating it into models of cognitive function) which opens new opportunities to advance biosocial paradigms.

## Materials and Methods

Using data from the Health and Retirement Study, genetic sample collection years 2006, 2008, 2010, we calculated and evaluated PGSs in 2,279 non-Hispanic Blacks and 9,991 non-Hispanic Whites. Further description of the HRS and our analysis sample can be found in supplemental materials (**S1**, **S2, Table S2**). We used a heavily modified R wrapper based on PRSice (6) and PLINK (5) to construct PGSs. We selected parameters reflecting the range of those used in the literature (e.g. (21, 22), **Table 1**). Detailed methods for selection of GWAS meta-analysis (**S3, Fig. S6**), mathematically removing a study from GWAS meta-analysis results (**S3**), PGSs estimation and assessment (**S4**), bootstrapping estimates of semi-partial R^2^ (**S4**), the overlap between SNPs across PGS estimation methods (**S5, Table S3**), a comparison of results for EA 2013 and EA 2016 (**S6, Fig. S7**), and a comparison of PGS results using the exact trait in a GWAS or an epidemiological definition (**S7**, **Fig. S8**) are described in the supplementary material.

## Acknowledgements

Funding for this work was supported by U01 AG009740, RC2 AG036495, RC4 AG039029. The authors would also like to thank the HRS participants for their contributions.

### Conflict of Interest

The authors declare no conflicts of interest.

